# The distinctive mechanical and structural signatures of residual force enhancement in myofibers

**DOI:** 10.1101/2023.02.19.529125

**Authors:** Anthony L. Hessel, Michel Kuehn, Bradley M. Palmer, Devin Nissen, Dhruv Mishra, Venus Joumaa, Johanna Freundt, Weikang Ma, Kiisa C. Nishikawa, Thomas Irving, Wolfgang A. Linke

## Abstract

In muscle, titin proteins connect myofilaments together and are thought to be critical for contraction, especially during residual force enhancement (RFE) when force is elevated after an active stretch. We investigated titin’s function during contraction using small-angle X-ray diffraction to track structural changes before and after 50% titin cleavage and in the RFE-deficient, *mdm* titin mutant. We report that the RFE state is structurally distinct from pure isometric contractions, with increased thick filament strain and decreased lattice spacing, most likely caused by elevated titin-based forces. Furthermore, no RFE structural state was detected in *mdm* muscle. We posit that decreased lattice spacing, increased thick filament stiffness, and increased non-crossbridge forces are the major contributors to RFE. We conclude that titin directly contributes to RFE.

**One-Sentence Summary:** Titin contributes to active force production and residual force enhancement in skeletal muscles.

## Main Text

Animal movement through complex environments requires unique mechanical features of not only isometric or shortening (concentric) muscle contractions, but also lengthening (eccentric) contractions (*1*). Eccentric contraction produces immediate and rapid force enhancement at the level of the sarcomere that is greater than that possible during isometric or concentric contractions and is poorly described by current muscle models (*2, 3*). Force enhancement improves functional tasks such as counter-movement jumps (*4*), downhill braking (*5*), and joint stabilization when negotiating complex terrain (*6*). Clinically, eccentric-focused training accelerates muscle hypertrophy above conventional training, and is widely used for athletes and also for restoring mobility independence to older adults (*7, 8*).

During eccentric contraction, both crossbridge and non-crossbridge structures are stretched and produce viscoelastic forces that contribute to force enhancement (*9*). It has been suggested that crossbridge-based force enhancement plateaus after a short stretch of ∼28 nm per sarcomere (*9, 10*), while further stretch increases force via non-crossbridge, viscoelastic “spring” elements that increase stiffness upon activation (*9, 11, 12*). After stretch, force enhancement from crossbridges dissipates quickly while the non-crossbridge elastic component remains, leaving a long-lasting enhancement of force above that of pure isometric contractions at the same final length, the so-called residual force enhancement (RFE)(*13*). The mechanisms underlying RFE are unclear but mounting evidence suggests that the titin protein plays a key role (Fig. 1A). Titin, the largest known protein, extends from the Z-disk to the M-line, where it runs along the thick filament in the A-band, and is extensible in the I-band. Titin extends during sarcomere stretch and becomes 4-6 times stiffer upon Ca^2+^ activation (*14*–*18*) via as yet unresolved mechanisms (*19*). Increasing titin-based forces during passive stretch (*20*) and submaximal activation (*21, 22*) also increases the Ca^2+^ sensitivity of force with increasing sarcomere length, implying that increased crossbridge recruitment may also contribute to RFE.

**Fig. 1.**
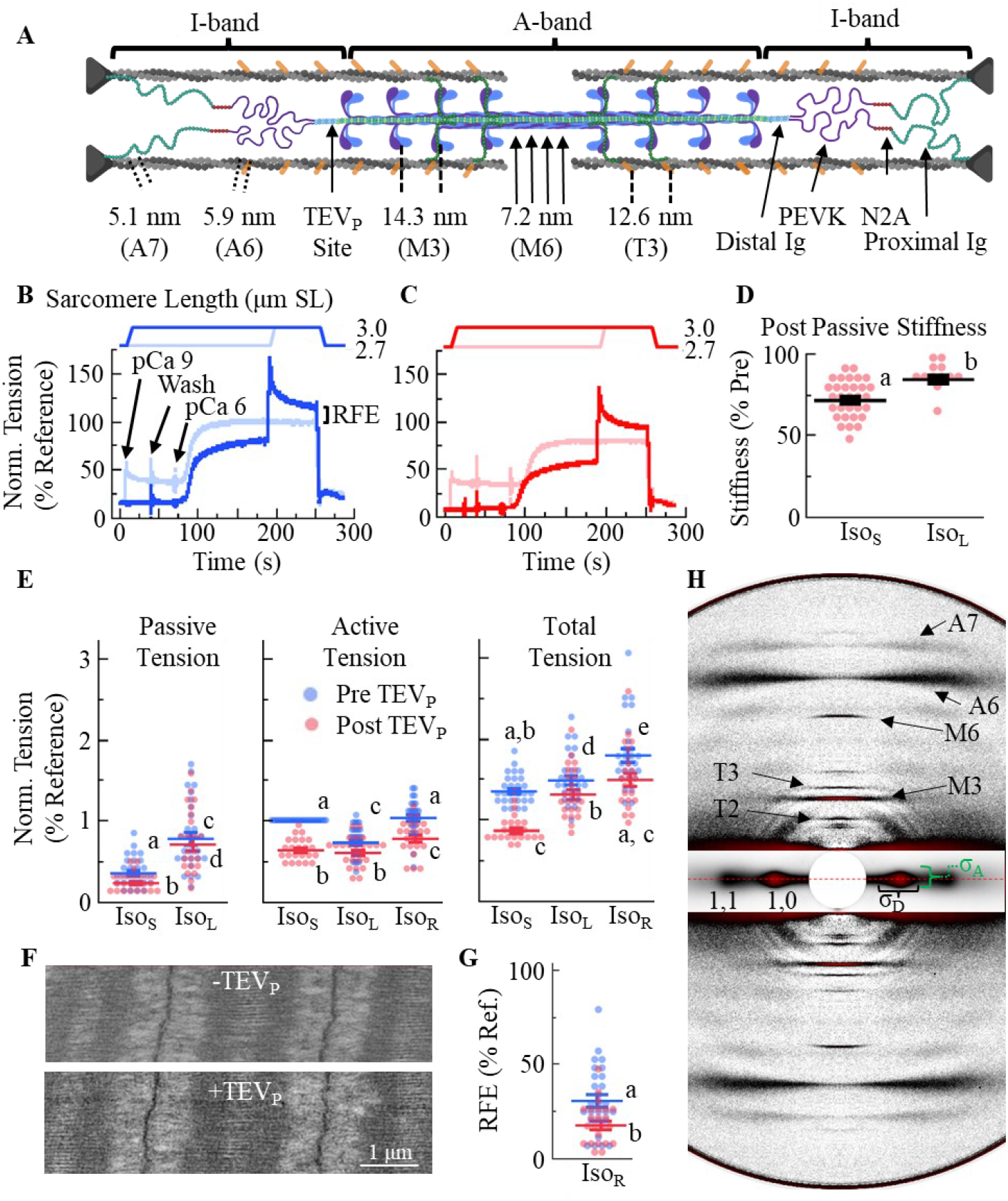
X-ray diffraction of TC fiber bundles before and 50% titin cleavage during Iso_S_, Iso_L_, and Iso_R_ conditions. (**A**) Schematic of skeletal half-sarcomere with relevant structural periodicities, I-band titin segments, and TEV_P_ cleavage site indicated. (**B-C**) Tension traces of fibers during mechanical experiments before (B) and after (C) 50% titin cleavage. Active tension is total tension during contractions, minus the passive tension at those lengths. (**D**) Passive stiffness after titin cleavage, normalized to its paired pre-stiffness value at Iso_S._ (**E**) Tension before (blue) and after (red) titin cleavage for passive, active, and total conditions. (**F**) transmission electron micrographs of TC sarcomeres after full mechanical protocols with sham (- TEV_P_) or treatment (+TEV_P_) conditions. (**G**) RFE in fibers before and after titin treatment. (**H**) Representative X-ray diffraction pattern of skeletal psoas fibers, with labeled reflections indicating relevant periodic structures that are referenced in (A). Connecting letters: different letters are significantly different (post hoc analysis P < 0.05). Data throughout reported as mean ± s.e.m. Full statistical details in Table S1. letters are significantly different (post hoc analysis P < 0.05). Data throughout reported as mean ± s.e.m. Full statistical details in Table S1.

We investigated titin’s function during skeletal muscle contraction using small-angle X-ray fiber diffraction to track structural changes before and after cleavage of 50% of I-band titin in mouse fast-twitch muscle fibers, and in fibers from ‘muscular dystrophy with myositis’ (*mdm*) mice which lack force enhancement (*23*). Specific cleavage of I-band titin was achieved by using heterozygotes of a transgenic ‘titin cleavage’ (TC) mouse model (Fig. S1), in which titin is controllably cleaved via an embedded tobacco etch virus protease (TEV_P_) cleavage site (*17, 24*). *Mdm* carries a complex titin mutation that manifests as a small deletion in the titin gene (*25, 26*). Our results demonstrate that, during RFE, in addition to increased tension, there are structural-based X-ray diffraction signatures that are distinct from those of pure isometric contraction. Titin cleavage attenuates RFE tension, blunts RFE-associated diffraction signatures, and increases disorder in the myofilament lattice. We further report that when RFE is absent, as in *mdm* mice, the structural signature of RFE is also absent. Collectively, we posit that RFE is caused, in part, by decreased lattice spacing, stretched inter-myofilament bridge elements (e.g., titin, myosin-binding protein C [MyBPc]), and increased thick filament stiffness, with titin a major regulator of each.

## Results and Discussion

### Titin cleavage reduces mechanical stability

We conducted RFE experiments on permeabilized heterozygote TC psoas fibers before and after 50% titin cleavage via TEV_P_ incubation. X-ray diffraction patterns and force traces were obtained under three conditions: isometric contraction at 2.7 μm sarcomere length (SL) (short isometric contraction; Iso_S_), isometric contraction at 3.0 μm SL (long isometric contraction; Iso_L_), and stretch-hold contraction from 2.7 to 3.0 μm SL (isometric contraction after an active stretch [RFE]; Iso_R_). Compared to pre-titin cleavage, post-cleavage fibers produced less tension, but the shape of the tension traces was similar (Fig. 1B-C). For Iso_S_ and Iso_L_, 50% titin cleavage reduced passive stiffness (measured here by sinusoidal oscillations; Fig. 1D), passive tension, active tension, and total tension (P < 0.01, Fig. 1E; Table S1), but the relative difference after treatment was always greater for Iso_S_ vs. Iso_L_ (Fig. S2A). Titin-based force and stiffness are known to be critical for thick filament centering in the A-band, with greater stability at longer SLs where titin-based forces are higher (*17, 27*). Indeed, at a near-slack SL of 2.4 μm, the A-bands of titin-cleaved fibers fell apart during maximal activation (pCa 4) (*17*), but were maintained here at longer SLs and at submaximal activation (pCa 6; Fig. 1F). Furthermore, RFE (increase in total force from Iso_L_ to Iso_R_) was reduced after titin cleavage (P < 0.01; Fig. 1G, S2B) from 30.69 ± 3.35% to 17.44 ± 2.24% of reference force (Iso_L_). We converted RFE to force per thick filament (f_TF_), assuming that all forces are transmitted to the thick filament, and that in samples with a d_1,0_ = 37.69 nm ∼80% of the cross-sectional area is occupied by myofibrils, resulting in a thick filament density of ∼488×10^6^ thick filaments/mm^2^. The active tension increased from 48.26 kPa in Iso_L_ to 68.53 kPa in Iso_R_, resulting in f_TF_ values of 98.95 pN and 140.51 pN for Iso_L_ and Iso_R_, respectively. Therefore, compared to Iso_L_, Iso_R_ increased by 41.56 pN. These results are consistent with numerous previous studies that provided indirect evidence for titin’s involvement in RFE (*28*–*30*), and with the absence of RFE in the *mdm* titin mutant (*31, 32*). These results demonstrate a direct relationship between I-band titin and RFE.

### Lattice structure and order during Iso_R_ are structurally distinct

We assessed structural changes using small-angle X-ray diffraction (Fig. 1H), which provides information concerning the underlying structural arrangements of sarcomeric proteins (Fig. 1A). We compared diffraction patterns from pure isometric contractions at two different sarcomere lengths (Iso_S_ vs. Iso_L_) and contractions with ramp stretch-hold from the short to the longer length (Iso_L_ vs Iso_R_). Lattice spacing (LS) was calculated as the separation of the 1,0 equatorial reflections from thick filaments (D_1,0_ plane; Fig 1H). Computational modeling studies suggest that increasing LS decreases force production and blunts length-dependent activation (LDA) by affecting crossbridge kinetics (*33*). LS is controlled in part by radial forces imposed on the myofilament lattice by titin (*34, 35*) and expands in relaxed fibers after titin cleavage (*22*). Furthermore, myopathic human fibers with abnormally stiff titins have smaller LS than controls (*36*). We report that LS decreased from Iso_S_ to Iso_L_, as expected, but decreased further in Iso_R_ (P< 0.0001; Fig. 2A, S3; Table S2). The “excess” lattice shrinkage in Iso_R_, noted previously (*37*), suggests that radial forces acting on the myofilament lattice are enhanced after active stretch, suggesting increased titin-based forces in Iso_R_ with respect to Iso_L_. LS is an important regulator of crossbridge kinetics and therefore cross-bridge based force production (*33*), especially in permeabilized preparations, where the lattice is generally expanded compared to intact, and shrinks more rapidly upon stretch. Compared to Iso_L_, the Iso_R_ lattice was smaller by 1.08% (0.41 nm). Based on previous assessments (*33*), we estimated a small but significant RFE contribution of ∼5.2% force, or f_TF_ ∼ 5.14 pN.

**Fig. 2.**
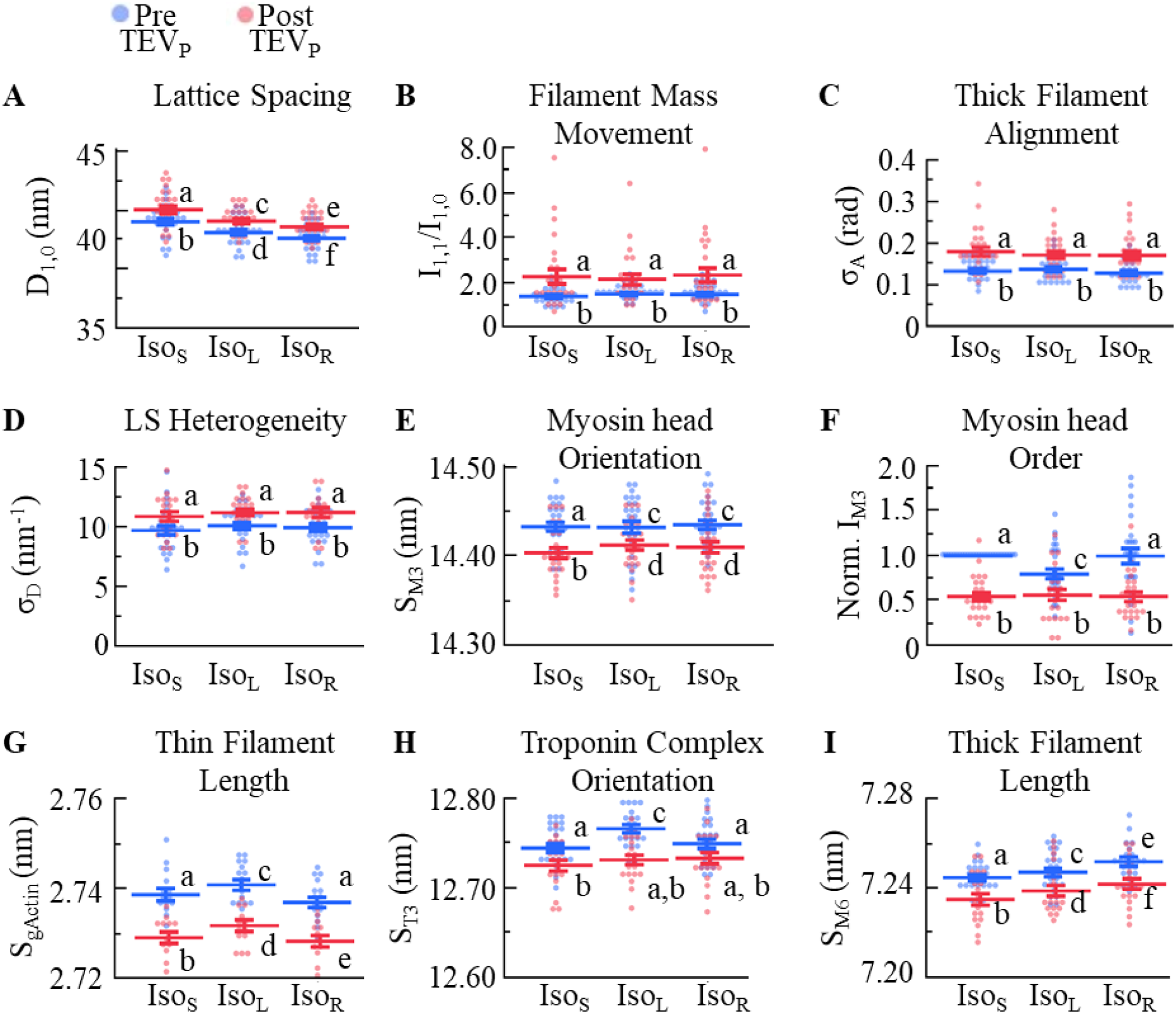
Sarcomeric structural parameters of TC fibers before and after 50% titin cleavage. D_1,0_ (**A**), I_1,1_/I_1,0_ (**B**), s_A_ (**C**), s_D_ (**D**) S_M3_ (**E**), I_M3_ (**F**), S_gActin_ (**G**), S_T3_ (**H**), and S_M6_ (**I**) were recorded before (blue) and after (red) 50% titin cleavage, at three conditions: Iso_S_, Iso_L_, and Iso_R_. Connecting letters: different letters are significantly different (Tukey HSD P < 0.05). Full statistical details in Table S2 and Table S3. Intra-sample pre-post differences in Fig. S3.

### Structural evidence of crossbridges suggest no differences between Iso_R_ and Iso_L_

We next considered whether more crossbridges are present during Iso_R_ vs. Iso_L_. Previous reports using structural, mechanical or ATP consumption assays suggested no differences in crossbridge recruitment between Iso_L_ and Iso_R_ (*9, 37*–*42*). The equatorial intensity ratio (I_1,1_/I_1,0_) is widely used as a measure of the transfer of mass (i.e., myosin heads) from the thick to thin filaments, with increasing I_1,1_/I_1,0_ indicating more myosin heads associated with the thin filaments (*43*). We found that before titin cleavage, I_1,1_/I_1,0_ remained consistent across contraction conditions (P = 0.18; Fig. 2B, S3;Table S2). At longer SLs, I_1,1_/I_1,0_ decreased as filament overlap decreased (*44*). Iso_L_ has less thick and thin filament overlap compared to Iso_S_, so I_1,1_/I_1,0_ would be expected to decrease if nothing else changed. Therefore, the finding that I_1,1_/I_1,0_ is not significantly different between short and long SL contraction conditions suggests a relatively larger degree of crossbridge recruitment at longer SLs (i.e., Iso_L_ and Iso_R_), as expected via myofilament LDA (*45*). These findings also imply that there is no further crossbridge recruitment in Iso_R_ vs. Iso_L_, a finding that corroborates other studies using different methodologies (*9, 38*–*40*).

Titin cleavage by 50% led to an overall increase in I_1,1_/I_1,0_ across contraction conditions, with a large increase in the spread of the values including some with unrealistically large values greater than the rigor state (2-3; *43*) (P < 0.0001; Fig. 2B; Table S2), which was initially puzzling. These results are likely due to increased lattice disorder after titin cleavage as evidenced by increased σ_A_, an indicator of thick filament disorder across adjacent myofibrils (*46*), and σ_D_, a measure of LS heterogeneity, after titin cleavage (P < 0.001; Fig. 2C-D; Table S2). Because we cannot easily uncouple the effects of lattice disorder (*44, 47*) and mass shift due to myosin head movement on I_1,1_/I_1,0_, it would be unwise to interpret increased I_1,1_/I_1,0_ after titin cleavage as indicating more transfer of myosin heads towards actin. However, in contrast to the equatorial reflections, meridional patterns are less affected by lattice disorder. Myosin head configuration can be evaluated via the spacing (S_M3_) and intensity (I_M3_) of the M3 myosin meridional reflection. Increases in S_M3_ are typically associated with increasing crossbridge recruitment, or the reorientation of myosin heads in a position that increases the chance of attachment (*48*). Before titin cleavage, S_M3_ varied among conditions as follows: Iso_S_ < Iso_L_ = Iso_R_ (P < 0.0001; Fig. 2E, S3; Table S2), similar to I_1,1_/I_1,0_ before titin cleavage, both suggesting increased crossbridge recruitment at longer SL, but unaffected by previous active stretch. In comparison, titin cleavage reduced S_M3_ across all conditions (P < 0.0001; Fig. 2E, S3; Table S2). Decreased S_M3_ across conditions after 50% titin cleavage suggests that titin is important in modulating myosin head action not only in passive muscle, but also during contraction—a point suggested in muscles from mice with titin mutations but never before shown by cutting of titin springs within a sample. Additional support for this notion comes from I_M3_, which provides additional details about variation in myosin head orientation. Before titin cleavage, I_M3_ decreased from Iso_S_ to Iso_L_ (P < 0.001; Fig. 2F; Table S3) as expected due to decreased filament overlap (*48, 49*). However, I_M3_ for Iso_R_ > Iso_L_ = Iso_S_ (P < 0.001; Fig. 2F), even though they are at the same SL, suggesting that the orientation of heads is more ordered in Iso_R_. Of note, after titin cleavage, I_M3_ was reduced in all conditions to similar values, suggesting that titin-based forces are partially responsible for myosin head order, and the conditional effect with passive or active stretch.

The thin filaments also provide structural information indicative of crossbridge recruitment (*50*). In passive cardiac and skeletal muscle, thin filaments extend and troponin complexes reorient with sarcomere stretch in a way that has been linked to increasing Ca^2+^ sensitivity (*20, 22*). The A6 (S_A6_) and A7 (S_A7_) spacings report on the left- and right-handed actin helical structures within the thin filament (Fig. S4;Table S3) and are used here to calculate the axial spacing of the actin monomers (S_gActin_) (*51, 52*), where S_gActin_ can be used as a measure of thin filament extension (*53, 54*). Also relevant for this discussion are structural changes in the troponin complex, indicated by changes in the spacing (S_T3_) of the third troponin meridional reflection T3 arising from the troponin complexes spaced every ∼37 nm along the thin filaments. We found that before titin cleavage, both S_gActin_ and S_T3_ are Iso_S_ < Iso_L_ = Iso_R_ (P < 0.01; Fig. 2G-H; Table S3). Iso_L_ = Iso_R_ is noteworthy because in purely isometric or passive conditions, the thin filament acts as a stiff Hookean spring so that changes in thin filament strain can be directly interpreted as changes in the total (passive + active) force exerted on the filament (*22, 50*), as observed from Iso_S_ to Iso_L_; Fig. 1E). However, while Iso_R_ forces are larger than Iso_L_, the thin filament strains are equal. This suggests that something distinct is occurring during eccentric contraction that decouples force and thin filament strain and may suggest that crossbridge-based force is decreased while non-crossbridge based force is increased. Another interesting result is that the S_T3_ is altered by thin filament length, as expected for a thin filament associated protein. However, ANCOVA showed that S_T3_ decreases independently of S_gActin_ (P < 0.001; Fig. 2H; Table S3), although the physiological relevance of this independent S_T3_ action is unknown.

Increased titin-based forces may not necessarily result in increased thin filament strain because titin’s tether points are far away from the tip of the thin filament; titin’s permanent interaction with the thin filament is close to the Z-disk (*55, 56*), and other potential alternative interaction sites are at or around the N2A region in the I-band (*57, 58*). Although a definitive mechanism remains elusive, another possibility is disruption of MyBPc thick-thin bridges during the eccentric phase of the Iso_R_ condition. MyBPc bridges are purported to be important for contraction (*22, 59*–*61*) and seem to be forcibly ruptured by a quick passive stretch (*22*). Therefore, a role for MyBPc to explain the Iso_R_ seems plausible and could be directly studied using an inducible “cut and paste” MyBPc mouse line (*62*).

### Thick filament strain suggests titin-based forces are enhanced in Iso_R_ vs. Iso_L_

Arguably the most contentious debate in the RFE field is whether titin contributes to RFE by producing more force during Iso_R_ vs. Iso_L_. The hypothesis is as follows: upon activation, titin stiffness increases by 4-6 times (*15, 17, 18, 39*), which would imply that during and after eccentric contractions, titin-based forces would be greater than if just activated at the longer SL. Mechanically, titin-based force pulls on and stretches the thick filament, with increasing titin force extending the thick filament (*21, 22, 34, 63*). We found that thick filament strain, quantified via the spacing of the M6 meridional reflection (S_M6_), increased progressively as follows: Iso_S_ < Iso_L_ < Iso_R_ (P < 0.0001; Fig. 2I, Table S2), suggesting higher titin-based force in Iso_R_ vs. Iso_L_. Additionally, 50% titin cleavage slightly reduced S_M6_ across all conditions (P < 0.0001 for both; Fig. 2I; Table S2) but the general relationship among conditions was preserved. These data indicate titin as a key contributor to thick filament strain.

In addition to titin, MyBPc can also apply a pulling force to strain the thick filament. To date, MyBPc plays an unclear role during contraction, but can also function like a spring and store some level of force (*61*), so storing force during an eccentric stretch seems plausible. On the other hand, MyBPc is relatively short and so would most likely rupture and reattach to the thin filament during the ∼150 nm per half thick filament eccentric stretch conducted here, minimizing its contribution to force exerted on the thick filament during Iso_R_, although some contribution cannot be discounted. Apart from MyBPc, crossbridge forces during contraction also strain the thick filament (*63*). However, our analysis above, along with other mechanical and biochemical studies (*9, 38*–*40*), suggest no differences in the degree of crossbridge recruitment between Iso_L_ and Iso_R_ so that the contribution of crossbridge-based thick filament strain should be similar between Iso_L_ and Iso_R_. Based on our analysis, we postulate that the thick filament strain in Iso_R_ relative to Iso_L_ is primarily due to increased titin-based force after active stretch.

We next used the thick filament strain data to provide estimates of f_TF_ during Iso_R_, which also includes the LS-dependent effects described above. Compared to Iso_L_, the thick filament in Iso_R_ is strained 0.0694% more, with 41.56 pN more f_TF_. The excess f_TF_ is a product of both the forces acting to strain the thick filament, such as parallel elastic elements (e.g., titin and possibly MyBPc), and the changes in force that arise primarily from stiffening of the thick filaments when they are stretched (*64, 65*). Approximating changes to thick filament stiffness (Stiff_TF_) is a non-trivial task, as thick filament stiffness is nonlinear (*21*). To accomplish this, we derived f_TF_ as a function of percent thick filament elongation (e_p_) using active tension-thin filament strain data from fast twitch mouse muscle (*21*), and converted to f_TF_ - thick filament strain. The data were described well by a third-order polynomial: f_TF_ = 2.98*10^−8^e_p_ + 709e_p3_, simplified to f_TF_ = 709e_p3_, with the Stiff_TF_-strain relationship equal to the derivative of f_TF_, Stiff_TF_ = 2127e_p2_. We next calculated the e_p_ values for Iso_L_ and Iso_R_ from S_M6_ values as follows: passive S_M6_ at 3.0 μm SL is 7.205 nm (*21, 22, 34, 63*), and the Iso_L_ and Iso_R_ S_M6_ = 7.247 and 7.252 nm, respectively. Therefore, from the passive state, the thick filaments are strained 0.586% and 0.655% during Iso_L_ and Iso_R_, respectively. Plugging the e_P_ values into the Stiff_TF_-strain equation produces 729.7 and 912.9 pN/e_p_ for Iso_L_ and Iso_R_, respectively. Therefore, the thick filament stiffness increases by 25.1% from Iso_L_ to Iso_R,_ which also reflects an up to ∼25.1% increase in f_TF_ (*66*). However, the exact relationship between Stiff_TF_ and f_TF_ is somewhat less than linear, depending on the properties of thick filament compliance. Because the relationship is not yet well defined (*64, 65*), we used a conservative lower bound of ∼0.5 Stiff_TF_ to f_TF_ ratio. Said another way, ∼12.4 pN up to a maximum of ∼24.8 pN of the measured RFE per thick filament (f_TF_ ∼ 41.6 pN) can be associated with mechanical stiffening of the thick filament.

### A loss of RFE also leads to a loss of distinctive diffraction signatures during Iso_R_

As described above, we characterized distinctive diffraction signatures during Iso_R_, but an open question was whether this structural state is required for RFE. To assess this point, we utilized the muscular dystrophy with myositis (*mdm*) mouse, which to our knowledge is the only skeletal mouse model in which RFE can be induced in wild-type but not in HOM *mdm* (*23, 32*). We assessed whether the structural indicators of the RFE state were also missing in HOM skeletal muscles. Experiments were run similarly to the TC experiments, but at different SLs for the three contraction types: isometric 2.4 μm SL (*mdm* Iso_S_), isometric 3.2 μm SL (*mdm* Iso_L_), and RFE (*mdm* Iso_R_; stretch-hold from 2.4 to 3.2 μm SL). Compared to TC fibers before titin cleavage, WT *mdm* fibers showed similar diffraction signatures of RFE for LS, S_M6_, I_1,1_/I_1,0_, s_A_, S_M3_, and S_A6_ (Fig. 3A-F; Table S4). In contrast, *mdm* fibers showed no statistical difference in any of these parameters between Iso_R_ and Iso_L_ except for I_1,1_/I_1,0_. Here, as in the TC experiments after titin cleavage, HOM *mdm* produced unusually large I_1,1_/I_1,0_, with associated increases in σ_A_, suggesting reduced thick filament stability in HOM *mdm* fibers, and so inflated I_1,1_/I_1,0_ independent of myosin head mass movement (see discussion above). Based on these data, it is reasonable to propose that there is a causative relationship between the distinctive diffraction signature during Iso_R_ and the presence of RFE.

**Fig. 3.**
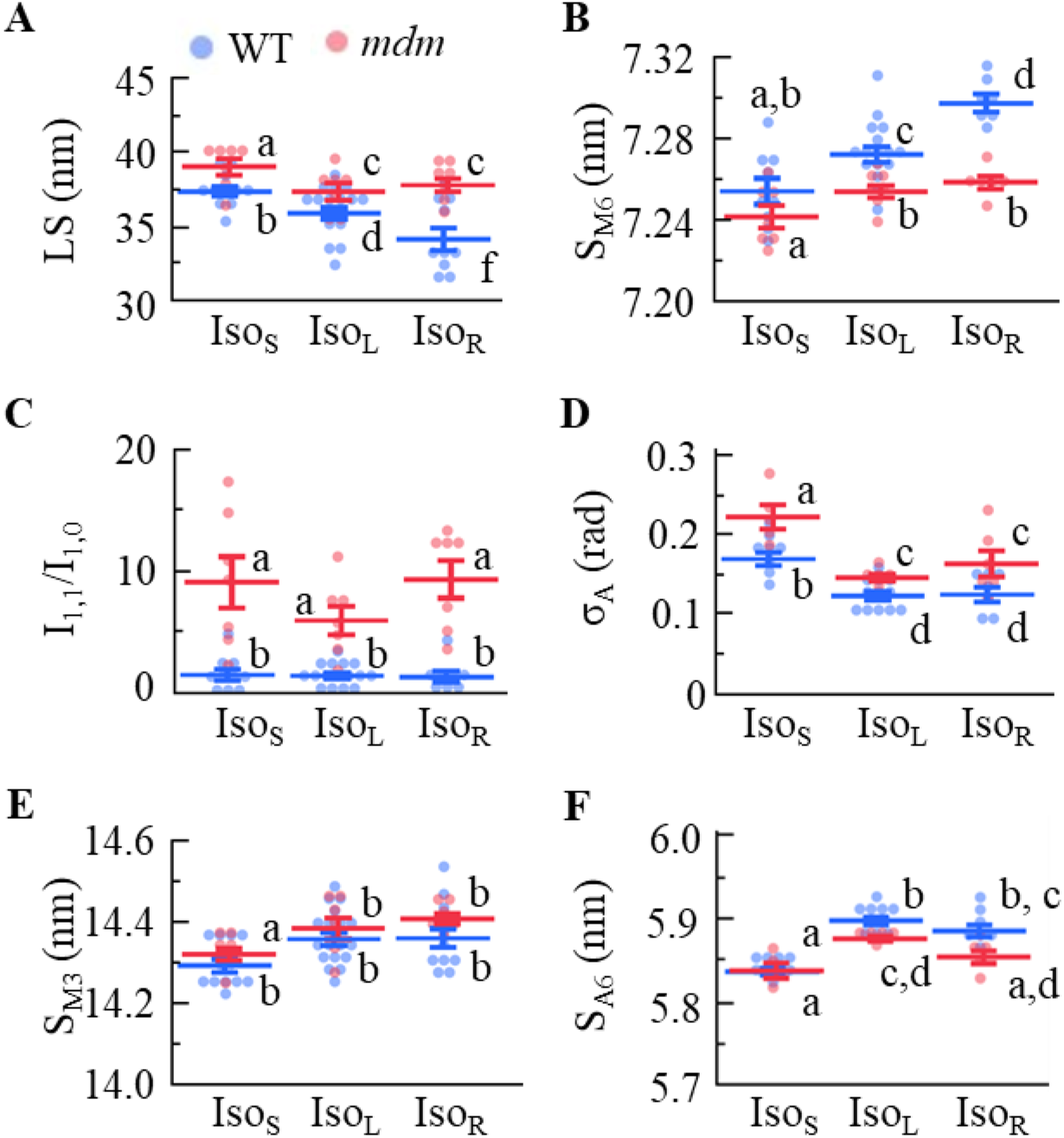
Titinopathic *mdm* fibers produce no distinctive RFE condition. D_1,0_ (**A**), S_M6_ (**B**), I_1,1_/I_1,0_ (**C**), s_A_ (**D**), S_M3_ (**E**), and S_A6_ (**F**) were recorded for WT (blue) and *mdm* (red) EDL fiber bundles at three conditions: Iso_S_, Iso_L_, and Iso_R_. Connecting letters: different letters are significantly different (Tukey HSD P < 0.05). Full statistical details in Table S4.

### A stiffer titin in Iso_R_ vs. Iso_L_ contributes to RFE

The underlying mechanism(s) of RFE, here f_TF_ = 41.6 pN above Iso_L_ tension, have long been unresolved. With this X-ray diffraction dataset, we thus far estimated two mechanisms that contribute to RFE: shrinking LS and increased thick filament stiffness (f_TF_ ∼ 12.4-24.8 pN). This implies that there is still ∼16.8-29.2 pN of f_TF_ unaccounted for. This most likely comes from the stretch of non-crossbridge myofilament bridge proteins. Titin is thought to become stiffer upon activation, and we can calculate how much stiffer it would become if it did indeed account for the missing component. On this approximation, we assume that titin contributes all forces imposed on the thick filament and that there are 12 titins per thick filament (6 titins per half-thick filament), equating to ∼2.8-4.9 pN per titin. Based on previous data (*67*), passive titin-based forces are ∼2.5 and ∼5 pN per titin at 2.7 and 3.0 μm SL, respectively, equating to a 2.7 to 3.0 μm SL stretch stiffness (∼150 nm stretch per titin) of ∼0.016 pN/nm. If we add the excess titin-based forces to the 3.0 μm SL values assuming it is all the unaccounted for f_TF_, then the titin-based stiffness after the stretch would be ∼0.035-0.049 pN/nm, suggesting that upon activation, titin stiffness increased ∼2.2-3.1 times compared to the passive state. This value will be somewhat lower if MyBPc or other parallel elastic components are involved. Presently, no molecular mechanism for this activation-dependent increase in titin stiffness is agreed upon, although studies are ongoing (*19, 58*). One idea (*11, 19, 57*, thoroughly discussed in *68*) relies on evidence of relatively weak titin-thin filament interaction at the N2A and nearby PEVK region in passive muscle and that this binding is stronger in the presence of Ca^2+^. This attachment functionally shortens the titin free-length and allows only the stiffer PEVK region to extend with increasing SL. Fig. 4 presents hypothetical configurations that can explain our data, where titin free-lengths are extended more during active vs. passive stretch, and deserves further empirical study. At present and under an as-yet unclear mechanism, we provide strong evidence that titin-based forces are greater after an active stretch as compared to an isometric contraction at the longer length, leading to structural and mechanical changes that enhance force, which answers the decades-old question as to why RFE exists.

**Fig. 4.**
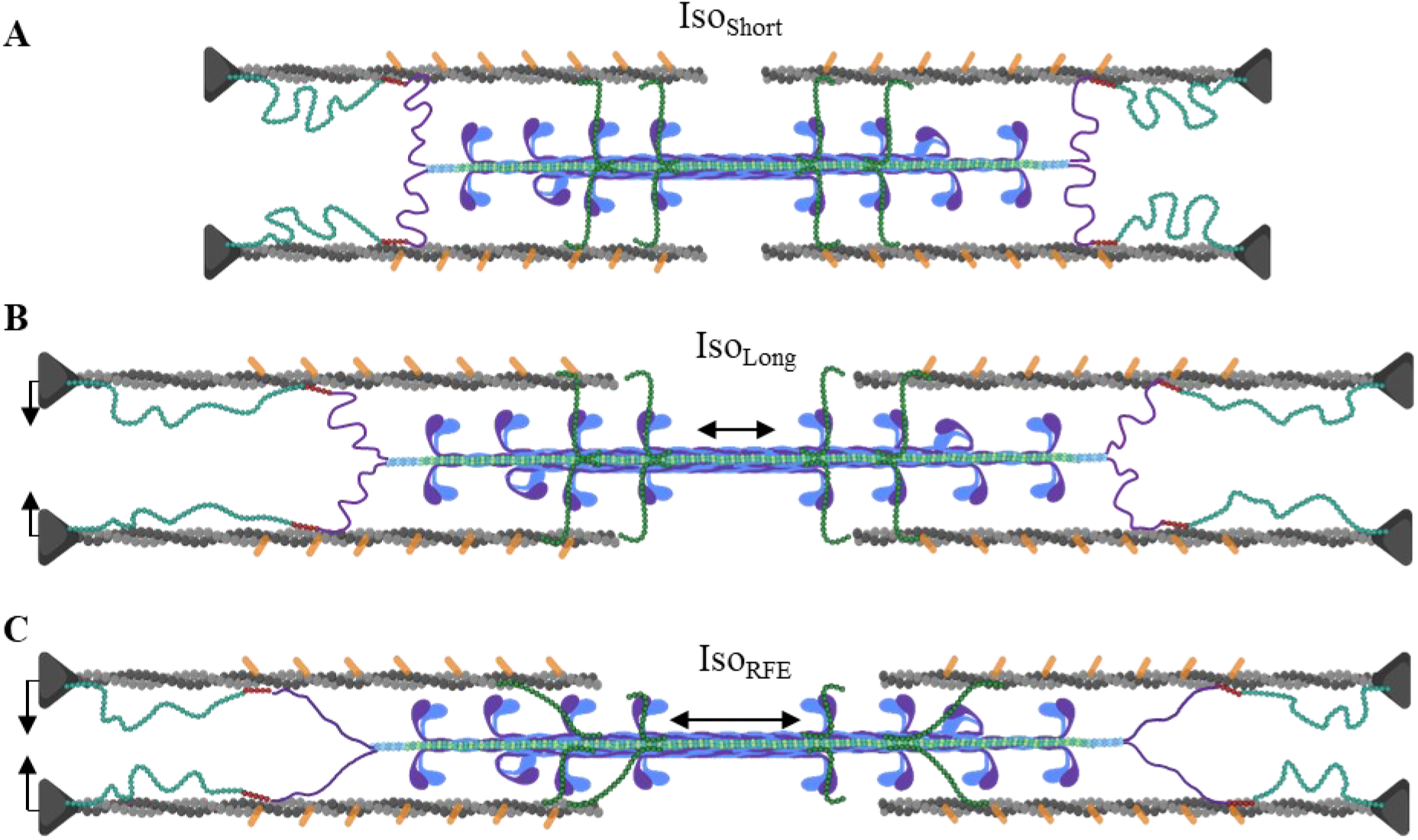
Mechanism of RFE. Configuration of sarcomeric proteins during (**A**) Iso_S_, (**B**) Iso_L_, and (**C**) Iso_R_ that can account for their distinctive mechanical and structural signatures. In passive muscle, low-level titin-thin filament interactions occur in such a way that passive stretch is enough to detach-reattach and/or drag titin along the thin filament so that titin-based free length and extension are still similar to if they were not attached at all. During contraction, the titin-thin filament interaction becomes stronger, so that during an eccentric contraction, titin extension occurs above that in passive, producing elevated titin-based force and explaining the mechanical and structural signatures in Iso_R_. Increased titin-based force contributes to RFE, which also leads to smaller lattice spacing and increased thick filament stiffness, improving force production and force transmission, respectively.

## Conclusions

Our data demonstrate that titin is a critical regulator of sarcomeric tension, and as such, an essential determinant of RFE. Furthermore, the presence of RFE seems to align with a distinctive structural state that is affected by titin cleavage and the *mdm* mutation. Finally, our analysis provides evidence that the generation of RFE is predominately caused by not only increases to titin-based forces, but also via titin’s pull on sarcomeric structures that decrease lattice spacing and increase thick filament stiffness.

## Acknowledgments

We thank the BioCAT beamline support staff at the APS for their steadfast commitment to our project, Andreas Unger for TEM imaging assistance, Beth Dennison for elite technical assistance, and Anna Good for critical text and artistic editing. This research used resources of the Advanced Photon Source, a U.S. Department of Energy (DOE) Office of Science User Facility operated for the DOE Office of Science by Argonne National Laboratory under Contract No. DE - AC02 - 06CH11357, and further NIH support. The content is solely the responsibility of the authors and does not necessarily reflect the official views of the National Institute of General Medical Sciences or the National Institutes of Health.

## Funding

German Research Foundation grant 454867250 (ALH)

German Research Foundation grant SFB1002A08 (WAL)

IZKF Münster Li1/029/20 (WAL)

National Institutes of Health P41 GM103622 (TI)

National Institute of Health P30 GM138395 (TI)

US National Science Foundation IOS-2016049 (KN)

## Author contributions

Conceptualization: ALH, WAL, KN

Methodology: ALH, WM, BP, BD, VJ, KN

Investigation: ALH, MK, WM, DN, BD, JF, DM, VJ, KN, BP

Visualization: ALH, MK, DN, BP

Funding acquisition: ALH, WAL, TI, KN

Project administration: ALH

Supervision: ALH, WAL

Writing – original draft: ALH

Writing – review & editing: All authors

## Data and materials availability

All data are available in the main text or the supplementary materials, or available upon reasonable request.

## Supplementary Materials

Materials and Methods

Supplementary Text

Figs. S1 to S4

Tables S1 to S4

References (*69*–*72*)

Data S1 (Separate file: Source_Data)

