## Supplemental Information for "The distinctive mechanical and structural signatures of residual force enhancement in myofibers"

Science
AAAS

### Supplementary Materials for

#### The distinctive mechanical and structural signatures of residual force enhancement in myofibers

**Authors:** Anthony L. Hessel[1*], Michel Kuehn[1], Bradley M. Palmer[2], Devin Nissen[3], Dhruv Mishra[4], Venus Joumaa[5], Johanna Freundt[1], Weikang Ma[3], Kiisa C. Nishikawa[4], Thomas Irving[3], Wolfgang A. Linke[1]

**This PDF file includes:**

Materials and Methods
Supplementary Text
Figs. S1 to S4
Tables S1 to S4
Captions for Data S1

**Other Supplementary Materials for this manuscript include the following:**

Data S1 Source_Data (separate excel file)

#### Materials and Methods

Animal model and muscle preparation

Titin cleavage (TC) mice. — Animal procedures were performed according to the guidelines of the local animal care and use committee and approved by the local authorities (LANUV NRW, 81-02.04.2019.A472). HaloTag-TEV (titin cleavage, TC, (*1*)) mice were bred and housed at the University Clinic Muenster. Genotyping was conducted by PCR in duplicate using custom primers: 5'cgtggtggcttatcttctagc3', 5'ctgttggttcatgcatctcc3', as previously described (*2*). Genetically heterozygous adult TC mice (age range, 2 – 6 months) were humanly euthanized and psoas muscle immediately extracted for long-term storage and permeabilized ("skinned) at -20°C using standard glycerol techniques (1:1 rigor : glycerol; rigor contains (mM) KCl (100), $MgCl_2$ (2), ethyleneglycol- bis(β-aminoethyl)-N,N,N′,N′-tetraacetic acid (EGTA,5), Tris (10), dideoxythymidin (DTT, 1), protease inhibitors [Complete, Roche Diagnostics, Mannheim, Germany], pH 7.0). Samples were shipped to the BioCAT facility on ice for all experimental tests and stored at -20°C until used. On the day of experiments, psoas muscles were removed from the storage solution and vigorously washed in relaxing solution (composition (in mM): potassium propionate (45.3), N,N-Bis(2-hydroxyethyl)-2-aminoethanesulfonic acid BES (40); EGTA (10), $MgCl_2$ (6.3), Na-ATP (6.1), DTT (10), protease inhibitors (complete), pH 7.0)). Bundles containing 15-30 fibers (3-6 mm long) were carefully excised and kept in physiological register by tying silk suture knots (sizing 6-0 or 4-0) at the distal and proximal ends of the bundle. Samples were then immediately transferred to the experimental chamber (see below).

Muscular dystrophy with myositis (*mdm*) mice. — The Institutional Animal Care and Use Committees at Northern Arizona University and Illinois Institute of Technology approved all husbandry and experimental protocols. Heterozygous mice of the strain B6C3Fe a/a-Ttn *mdm*/J were obtained from the Jackson Laboratory (Bar Harbor, ME, USA) and a breeding colony was maintained to produce wild type and homozygous recessive mice (*mdm*). Wild type and *mdm* mice were sacrificed at 24-30 days via isoflurane gas overdose confirmed by cervical dislocation. Extensor digitorum longus (EDL) muscles were extracted from euthanized mice following (*3*). From here, the protocol is the same as in TC mice.

Small angle X-ray diffraction and fiber mechanics apparatus

X-ray diffraction patterns were collected using the small-angle instrument on the BioCAT beamline 18ID at the Advanced Photon Source, Argonne National Laboratory (*4*). The X-ray beam (0.103 nm wavelength) was focused to ~0.06 x 0.15 mm at the detector plane, with an incident flux of ~$3x10^{12}$ photons per second. The sample to detector distance was set between 2.0 and 3.5 m, and the X-ray fiber diffraction patterns collected with a downstream CCD-based X-ray detector (TC experiments: Mar 165, Rayonix Inc, USA; *mdm* experiments: PCCD 16080; Aviex, New York, USA). For TC experiments, diffraction patterns were captured with 1 s exposure times, while for *mdm* experiments, a series of 25 or 50 X-ray photographs (10 ms X-ray exposure per photo over 1 s) were collected and combined. An inline camera built into the system allowed for initial alignment with the X-ray beam and continuous sample visualization during the experiment. Muscle preps were hung on custom muscle mechanics rigs, as explained previously (*5*, *6*). Sarcomere length (SL) was measured via laser diffraction using a 4-mW Helium-Neon laser. Force baseline was set at slack length. After this initial setup, fiber length changes were accomplished through computer control of the motor. Experiments were conducted at 25°C. The mechanical rig was supported on a custom designed motorized platform that allowed placement of muscle into the X-ray flight path and small movements to target X-ray exposure during experiments. Using the inline camera of the X-ray apparatus, the platform was moved to target the beam at different locations along the length of the sample. To limit X-ray exposure of any one part of the preparation, no part of the sample was exposed more than once. Fiber diameter was measured using the inline camera, and physiological cross-sectional area calculated at initial fiber length, with the assumption that the sample was a uniform cylinder longitudinally.

Experimental protocols and analysis

For TC experiments, sarcomere length (SL) was set to an initial length (~2.7 µm SL; $L_0$). Length changes were accomplished by manual or computer-driven means. After attachment, fibers underwent several cycles of rapid sinusoidal oscillations (5 x 30s, with 30s rest; 50 Hz, ± 5% $L_0$) on the relaxed preparation to acclimate the sample to the testing environment and detach any leftover cellular debris (e.g., collagen, vasculature) that would otherwise impact force measurements. Passive forces typically stabilized after 2-3 trials. The last oscillation series was used to quantify passive muscle stiffness, defined here as the averaged minimum-to-maximum force change of the last 10 oscillations, divided by the stretch amplitude.

The main mechanical tests consisted of two activation protocols, conducted in random order. For the first protocol, the relaxed fiber bundle was stretched from 2.7 → 3.0 µm SL at 1.0 µm SL $s^{-1}$ and held at that length for 30 s to allow for stretch relaxation. The sample was then moved into a bath of washing solution for 30 seconds x three washes each, followed by a transfer into activating solution x 2 washes (pCa = 6.0). Active forces rose until reaching a plateaued, was maintained for up to 2 minutes, and then shortened back to 2.7 µm SL and deactivated via 2 solution exchanges with relaxing solution. For protocol 2, the relaxed sample was kept at 2.7 µm SL and transferred into washing for 30 seconds x 3 exchanges, and then activating solution x 2 exchanges. When active forces reached a plateau, an eccentric stretch was performed from 2.7 → 3.0 µm SL at 1.0 µm SL $s^{-1}$ and held at that length for 60 s to allow for stretch relaxation. The preparation was then shortened back to 2.7 µm SL and transferred into relaxing solution x 2 exchanges. For both protocols, isometric active contraction times were adjusted to be as equal as possible to allow for a fare comparison between the protocols. The order of the two protocols was randomly assigned to each sample. Following the two protocols, the sample was incubated with $TEV_P$ for 20 mins (100 units $acTEV_P$ in 300 µl relaxing solution). After incubation, fibers were rinsed in fresh relaxing solution to remove excess protease, and several sinusoidal oscillations (see above) performed to measure stiffness. X-ray diffraction patterns were collected during isometric contraction conditions at 2.7 and 3.0 µm SL, and after the stretch-hold (RFE) condition. *Mdm* experiments were conducted similarly, but with short and long lengths of 2.4 and 3.2 µm SL, respectively.

X-ray image analysis

X-ray diffraction patterns were analyzed using the MuscleX open-source data reduction package (*7*). The "Scanning Diffraction" routine was used to measure the angular divergence of the 1,0 equatorial reflection. The routine obtains 2D and 1D radially integrated intensities of the equatorial intensities, and then fits Gaussian functions over the diffraction peaks to calculate the standard deviation (width σ) intensity distribution pattern. In this process, the routine obtains the integrated intensity of each equatorial reflection as a function of the integration angle. Gaussian profiles are fit to the projected peak intensities to find the standard deviation of the orientation angle (angle σ; $\sigma_A$) as a measure of the angular divergence of the angle that the sarcomeres in the myofibrils make to the long axis of the preparation. $\sigma_A$ is used as a proxy for inter-thick filament ordering (*8*). The "Equator" routine of Muscle X was used to calculate the $I_{1,1}$ / $I_{1,0}$ intensity ratio (IR), lattice spacing (LS) between thick filaments, and $\sigma_D$, a measure of the variability in thick filament lattice spacing (a proxy for lattice ordering). Meridional ($S_{T2}$, $S_{M3}$, $I_{M3}$, $S_{T3}$, $S_{M6}$) and off-meridional reflections ($S_{A6}$, $S_{A7}$) were collected using the MuscleX routines "Diffraction Centroids" and "Projection Traces". Actin monomer spacing ($S_{gActin}$) was calculated using A6 and A7 spacing data in an equation reported previously (*9*). Every image provides reflections of different quality, which lead to various levels of Gaussian fit errors for each reflection modeled, which increases the variation in spacings in the dataset. To limit these effects, fit errors > 10% were discarded. Positions of X-ray reflections on the diffraction patterns in pixels were converted to sample periodicities in nm using the 100-diffraction ring of silver behenate at $d_{001}$ = 5.8380 nm.

Statistics

Statistical analysis was conducted using JMP Pro (V16, SAS Institute Inc., Cary, NC, USA). Significance level was $\alpha = 0.05$. Response variables included all X-ray parameters. We first built a repeated-measures analysis of variance (ANOVA) design. For TC experiments, we used fixed effects treatment (pre /post $TEV_P$ incubation), and condition ($Iso_{Short}$, $Iso_{Long}$, $Iso_{RFE}$), and a treatment x condition interaction term, and a random (repeated-measures) effect of individual. For *mdm* experiments, we used fixed effects genotype (WT / *mdm*), and condition ($Iso_{Short}$, $Iso_{Long}$, $Iso_{RFE}$), and a genotype x condition interaction term, and a random (repeated-measures) effect of individual nested within genotype. Data was best Box-Cox transformed to meet assumptions of normality and homoscedacity when necessary, which were assessed by residual analysis, Shapiro-Wilk's test for normality, and Levene's test for unequal variance. Significant main effects were subject to Tukey's highly significant difference (HSD) multiple comparison procedures to assess differences between factor levels. This data is indicated in graphs via so-called connecting letters, where factor levels sharing a common letter are not significantly different from each other.

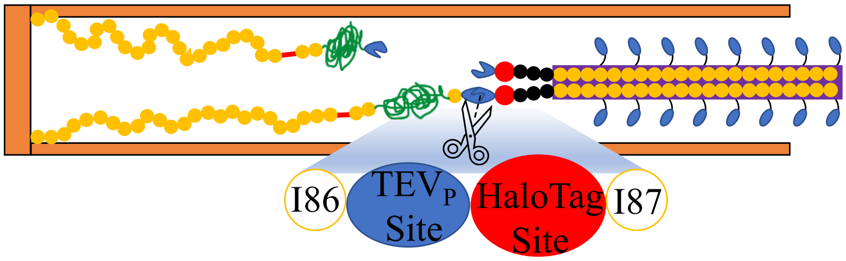

**Fig. S1. Titin cleavage (TC) mice allowed for intra-sample assessments of titin function.** A clean experimental strategy to study titin function is to specifically cleave I-band titin and terminate its functionality in a targeted and controllable fashion, allowing for changes to be tracked within the same preparation. The titin cleavage (TC) mouse model was generated with a cloned-in HaloTag-TEV cassette inserted into I-band titin close to the A-band (*24*). The Tobacco Etch Virus (TEV) protease recognition site is specifically cleaved by the TEV protease, while the HaloTag domain allows for easy protein labeling, useful for the assessment of titin cleaving. The insertion itself does not affect mouse development, muscle structure, or performance (*17*, *24*) and so allows for the study of as purely a titin-based effect as possible. Figure used with permission from Hessel et al., 2022).

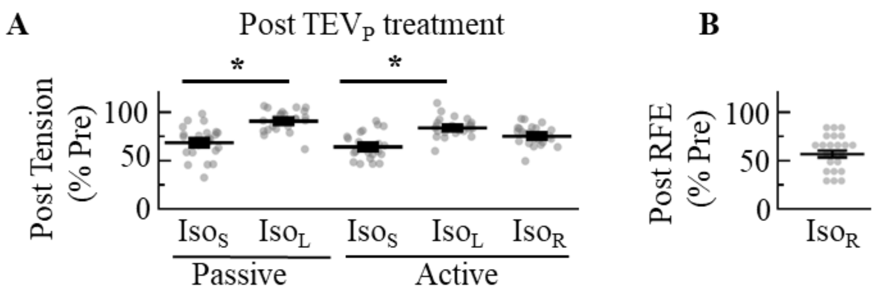

**Fig. S2. Tension differences after 50% titin cleavage from mechanical experiments.**

(**A**) The post tension values normalized to the paired pre-values for the passive component and active components during $Iso_S$, $Iso_L$, $Iso_R$ (defined in main text; separate data shown in Fig. 1E). (**B**) The post RFE values normalized to the paired pre RFE values (separate values shown in Fig. 1G) * $P < 0.05$ between the short ($Iso_S$) and long ($Iso_L$) conditions, assessed via ANOVA. Data shown as mean ± s.e.m.

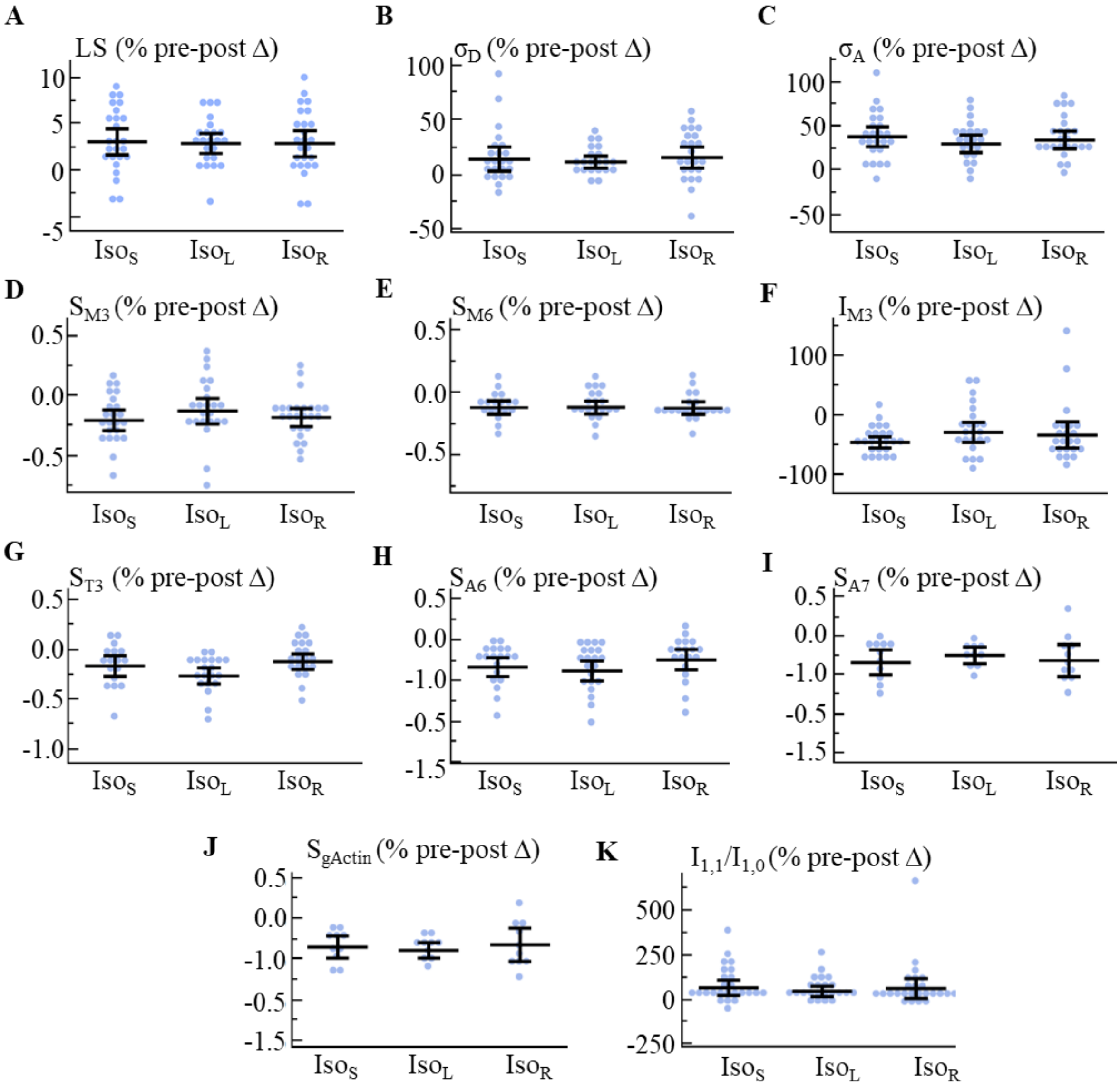

**Fig. S3. Differences in parameters recorded by X-ray diffraction from mechanical experiments after 50% titin cleavage.** (**A-K**) The post values for each X-ray diffraction parameter normalized to the paired pre-values. Data shown as mean ± 95% confidence interval of the mean.

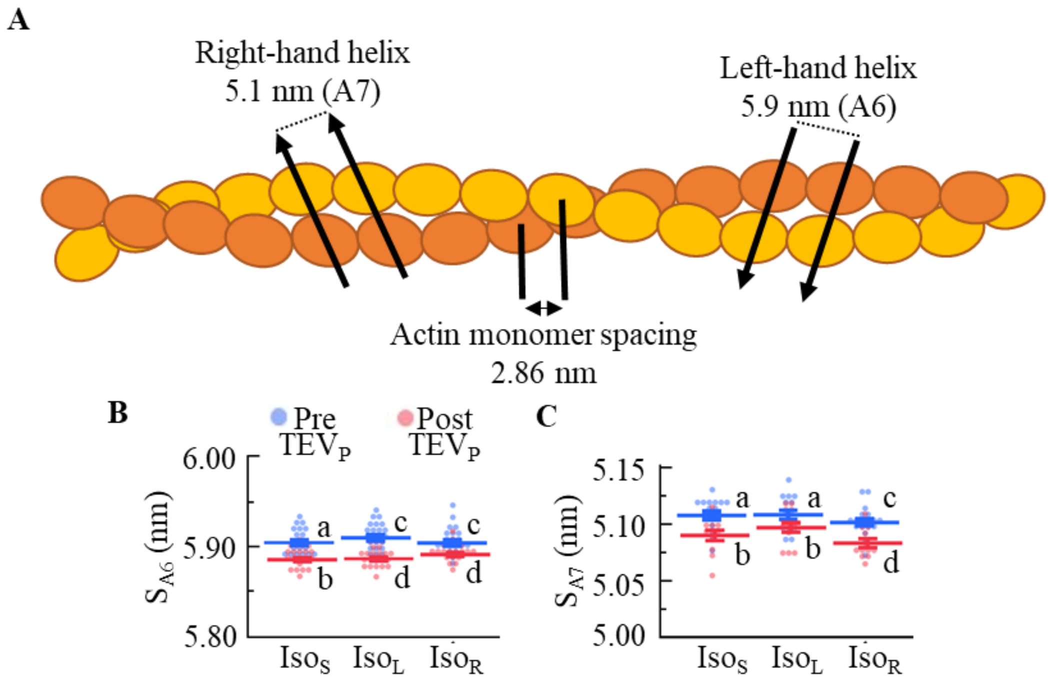

**Fig. S4. Thin filament strain. (A)** The actin double helix provides two repeating structures that are resolvable in our diffraction patterns, the left-handed helix (~5.9 nm repeat; $S_{A7}$), and the right-hand helix (~5.1 nm repeat; $S_{A6}$). The axial spacing of individual actin monomers ($S_{gActin}$, 2.86 nm) can be estimated with an equation (see methods). We provide spacing data for $S_{A6}$ (**B**) and $S_{A7}$ (**C**). The resulting actin monomer spacing, $S_{gActin}$ is presented in Fig. 2G. Connecting letters: different letters are significantly different (Tukey HSD $P < 0.05$). Data shown as mean ± s.e.m. Statistical details in Table S3.

**Table S1. Mechanics dataset, before and after TEV protease treatment (% Pre $Iso_{Short}$), from data in Figure 1. The ANOVA analysis F-stats and P-values are provided, as well as a connecting letter report from a Tukey's HSD analysis. Data reported as mean ± s.e.m. *Significant (P < 0.05).**

| Parameter (n) | Condition | Mean ± s.e.m. | ANOVA Main Effects | F | P | Connecting Letters |
|---|---|---|---|---|---|---|
| **Passive Tension (28)** | Pre $Iso_{Short}$ | 0.35 ± 0.03 | Treatment | 7.22 | 0.009* | a |
| **Passive Tension (24)** | Post $Iso_{Short}$ | 0.23 ± 0.02 | Condition | 159.11 | <0.0001* | b |
| **Passive Tension (28)** | Pre $Iso_{Long}$ | 0.78 ± 0.08 | Interaction | 0.46 | 0.46 | c |
| **Passive Tension (24)** | Post $Iso_{Long}$ | 0.71 ± 0.08 | | | | d |
| **Active Tension (28)** | Pre $Iso_{Short}$ | 1 | Treatment | 160.09 | <0.0001* | a |
| **Active Tension (23)** | Post $Iso_{Short}$ | 0.64 ± 0.03 | Condition | 54.72 | <0.0001* | b |
| **Active Tension (27)** | Pre $Iso_{Long}$ | 0.73 ± 0.03 | Interaction | 13.33 | <0.0001* | c |
| **Active Tension (24)** | Post $Iso_{Long}$ | 0.61 ± 0.04 | | | | b |
| **Active Tension (27)** | Pre $Iso_{RFE}$ | 1.04 ±0.04 | | | | a |
| **Active Tension (23)** | Post $Iso_{RFE}$ | 0.78 ± 0.04 | | | | c |
| **Total Tension (28)** | Pre $Iso_{Short}$ | 1.35 ± 0.03 | Treatment | 145.70 | <0.0001* | a,b |
| **Total Tension (23)** | Post $Iso_{Short}$ | 0.87 ± 0.03 | Condition | 131.82 | <0.0001* | c |
| **Total Tension (27)** | Pre $Iso_{Long}$ | 1.49 ± 0.06 | Interaction | 10.90 | <0.0001* | d |
| **Total Tension (24)** | Post $Iso_{Long}$ | 1.31 ± 0.06 | | | | b |
| **Total Tension (27)** | Pre $Iso_{RFE}$ | 1.80 ± 0.08 | | | | e |
| **Total Tension (23)** | Post $Iso_{RFE}$ | 1.49 ± 0.08 | | | | a,c |
| **Passive Stiffness (31)** | Post $Iso_{Short}$ | 0.72 ± 0.02 | Condition | 13.41 | 0.0007* | a |
| **Passive Stiffness (12)** | Post $Iso_{Long}$ | 0.84 ± 0.02 | | | | b |

**Table S2. Changes in X-ray reflection data after TEV protease treatment, from data in Figure 2. The ANOVA analysis F-stats and P-values are provided, as well as a connecting letter report from a Tukey's HSD analysis. Data reported as mean ± s.e.m. * P < 0.05.**

| Feature (n) | Treatment, Condition | Mean ± s.e.m. | Main Effects | F | P | Connecting Letters |
|---|---|---|---|---|---|---|
| LS (27) | Pre $Iso_{Short}$ | 38.99 ± 0.26 | Treatment | 55.92 | <0.0001* | a |
| LS (25) | Post $Iso_{Short}$ | 40.02 ± 0.32 | Condition | 33.23 | <0.0001* | b |
| LS (26) | Pre $Iso_{Long}$ | 38.10 ± 0.22 | Interaction | 0.06 | 0.94 | c |
| LS (24) | Post $Iso_{Long}$ | 39.06 ± 0.20 | | | | d |
| LS (25) | Pre $Iso_{RFE}$ | 37.69 ± 0.21 | | | | e |
| LS (25) | Post $Iso_{RFE}$ | 38.57 ± 0.22 | | | | f |
| $\sigma_A$ (26) | Pre $Iso_{Short}$ | 0.13 ± 0.005 | Treatment | 183.13 | <0.0001* | a |
| $\sigma_A$ (24) | Post $Iso_{Short}$ | 0.18 ± 0.01 | Condition | 2.49 | 0.09 | b |
| $\sigma_A$ (25) | Pre $Iso_{Long}$ | 0.14 ± 0.01 | Interaction | 1.00 | 0.37 | a |
| $\sigma_A$ (23) | Post $Iso_{Long}$ | 0.17 ± 0.01 | | | | b |
| $\sigma_A$ (24) | Pre $Iso_{RFE}$ | 0.13 ± 0.01 | | | | a |
| $\sigma_A$ (24) | Post $Iso_{RFE}$ | 0.17 ± 0.01 | | | | b |
| $\sigma_D$ (25) | Pre $Iso_{Short}$ | 9.69 ± 0.37 | Treatment | 30.39 | <0.0001* | a |
| $\sigma_D$ (23) | Post $Iso_{Short}$ | 10.85 ± 0.41 | Condition | 1.20 | 0.30 | b |
| $\sigma_D$ (24) | Pre $Iso_{Long}$ | 10.07 ± 0.30 | Interaction | 0.13 | 0.88 | a |
| $\sigma_D$ (22) | Post $Iso_{Long}$ | 11.18 ± 0.26 | | | | b |
| $\sigma_D$ (23) | Pre $Iso_{RFE}$ | 9.87 ± 0.34 | | | | a |
| $\sigma_D$ (23) | Post $Iso_{RFE}$ | 11.19 ± 0.41 | | | | b |
| $I_{1,1}/I_{1,0}$ (27) | Pre $Iso_{Short}$ | 1.37 ± 0.07 | Treatment | 24.7364 | <0.0001* | a |
| $I_{1,1}/I_{1,0}$ (25) | Post $Iso_{Short}$ | 2.25 ± 0.33 | Condition | 1.7432 | 0.18 | b |
| $I_{1,1}/I_{1,0}$ (26) | Pre $Iso_{Long}$ | 1.47 ± 0.06 | Interaction | 0.0773 | 0.93 | a |
| $I_{1,1}/I_{1,0}$ (24) | Post $Iso_{Long}$ | 2.12 ± 0.24 | | | | b |
| $I_{1,1}/I_{1,0}$ (25) | Pre $Iso_{RFE}$ | 1.47 ± 0.07 | | | | a |
| $I_{1,1}/I_{1,0}$ (25) | Post $Iso_{RFE}$ | 2.32 ± 0.31 | | | | b |
| $S_{M3}$ (26) | Pre $Iso_{Short}$ | 14.43 ± 0.01 | Treatment | 60.671 | <0.0001* | a |
| $S_{M3}$ (24) | Post $Iso_{Short}$ | 14.40 ± 0.01 | Condition | 0.788 | 0.457 | b |
| $S_{M3}$ (26) | Pre $Iso_{Long}$ | 14.43 ± 0.01 | Interaction | 0.746 | 0.477 | c |
| $S_{M3}$ (24) | Post $Iso_{Long}$ | 14.41 ± 0.01 | | | | d |
| $S_{M3}$ (26) | Pre $Iso_{RFE}$ | 14.44 ± 0.01 | | | | c |
| $S_{M3}$ (25) | Post $Iso_{RFE}$ | 14.41 ± 0.01 | | | | d |
| $S_{M6}$ (22) | Pre $Iso_{Short}$ | 7.244 ± 0.001 | Treatment | 59.15 | <0.0001* | a |
| $S_{M6}$ (20) | Post $Iso_{Short}$ | 7.233 ± 0.003 | Condition | 12.02 | <0.0001* | b |
| $S_{M6}$ (25) | Pre $Iso_{Long}$ | 7.247 ± 0.002 | Interaction | 0.13 | 0.88 | c |
| $S_{M6}$ (21) | Post $Iso_{Long}$ | 7.239 ± 0.002 | | | | d |
| $S_{M6}$ (24) | Pre $Iso_{RFE}$ | 7.252 ± 0.002 | | | | e |
| $S_{M6}$ (20) | Post $Iso_{RFE}$ | 7.242 ± 0.002 | | | | f |

**Table S3. Changes in X-ray reflection data after TEV protease treatment, from data in Figure 2. The ANOVA analysis F-stats and P-values are provided, as well as a connecting letter report from a Tukey's HSD analysis. Data reported as mean ± s.e.m. * P < 0.05.**

| Feature (n) | Treatment, Condition | Mean ± s.e.m. | Main Effects | F | P | Connecting Letters |
|---|---|---|---|---|---|---|
| $S_{T3}$ (24) | Pre $Iso_{Short}$ | 12.74 ± 0.005 | Treatment | 52.55 | <0.0001* | a |
| $S_{T3}$ (19) | Post $Iso_{Short}$ | 12.73 ± 0.01 | Condition | 7.69 | 0.001* | b |
| $S_{T3}$ (23) | Pre $Iso_{Long}$ | 12.77 ± 0.004 | Interaction | 3.15 | 0.047* | c |
| $S_{T3}$ (21) | Post $Iso_{Long}$ | 12.73 ± 0.01 | | | | a, b |
| $S_{T3}$ (22) | Pre $Iso_{RFE}$ | 12.75 ± 0.01 | | | | a |
| $S_{T3}$ (20) | Post $Iso_{RFE}$ | 12.73 ± 0.01 | | | | a, b |
| $S_{A6}$ (19) | Pre $Iso_{Short}$ | 5.918 ± 0.002 | Treatment | 92.21 | <.0001 | a |
| $S_{A6}$ (21) | Post $Iso_{Short}$ | 5.900 ± 0.003 | Condition | 1.98 | 0.14 | b |
| $S_{A6}$ (23) | Pre $Iso_{Long}$ | 5.923 ± 0.002 | Interaction | 1.17 | 0.22 | c |
| $S_{A6}$ (20) | Post $Iso_{Long}$ | 5.902 ± 0.003 | | | | d |
| $S_{A6}$ (19) | Pre $Iso_{RFE}$ | 5.920 ± 0.003 | | | | c |
| $S_{A6}$ (20) | Post $Iso_{RFE}$ | 5.906 ± 0.004 | | | | d |
| $S_{A7}$ (15) | Pre $Iso_{Short}$ | 5.107 ± 0.004 | Treatment | 49.89 | <0.0001* | a |
| $S_{A7}$ (13) | Post $Iso_{Short}$ | 5.089 ± 0.004 | Condition | 9.79 | 0.0002* | b |
| $S_{A7}$ (15) | Pre $Iso_{Long}$ | 5.108 ± 0.004 | Interaction | 0.88 | 0.42 | a |
| $S_{A7}$ (14) | Post $Iso_{Long}$ | 5.097 ± 0.004 | | | | b |
| $S_{A7}$ (18) | Pre $Iso_{RFE}$ | 5.101 ± 0.003 | | | | c |
| $S_{A7}$ (13) | Post $Iso_{RFE}$ | 5.083 ± 0.004 | | | | d |
| $S_{gActin}$ (14) | Pre $Iso_{Short}$ | 2.738 ± 0.001 | Treatment | 99.92 | <0.0001* | a |
| $S_{gActin}$ (12) | Post $Iso_{Short}$ | 2.729 ± 0.001 | Condition | 5.75 | 0.005* | b |
| $S_{gActin}$ (15) | Pre $Iso_{Long}$ | 2.741 ± 0.001 | Interaction | 0.06 | 0.94 | c |
| $S_{gActin}$ (14) | Post $Iso_{Long}$ | 2.732 ± 0.001 | | | | d |
| $S_{gActin}$ (16) | Pre $Iso_{RFE}$ | 2.737 ± 0.001 | | | | a |
| $S_{gActin}$ (12) | Post $Iso_{RFE}$ | 2.728 ± 0.001 | | | | e |
| $I_{M3}$ (26) | Pre $Iso_{Short}$ | 1 | Treatment | 98.16 | <0.0001* | a |
| $I_{M3}$ (24) | Post $Iso_{Short}$ | 0.54 ± 0.04 | Condition | 2.43 | 0.093* | c |
| $I_{M3}$(26) | Pre $Iso_{Long}$ | 0.79 ± 0.05 | Interaction | 3.53 | 0.032* | b |
| $I_{M3}$ (24) | Post $Iso_{Long}$ | 0.56 ± 0.06 | | | | c |
| $I_{M3}$ (26) | Pre $Iso_{RFE}$ | 0.99 ± 0.09 | | | | a, b |
| $I_{M3}$ (24) | Post $Iso_{RFE}$ | 0.54 ± 0.04 | | | | c |

**Table S4. X-ray reflection data between WT and *mdm* muscle, from data in Figure 3. The ANOVA analysis F-stats and P-values are provided, as well as a connecting letter report from a Tukey's HSD analysis. Data reported as mean ± s.e.m. * P < 0.05.**

| Parameter (n) | Genotype, Condition | Mean ± s.e.m. | ANOVA Main Effects | F | P | Connecting Letters |
|---|---|---|---|---|---|---|
| LS (11) | WT $Iso_{Short}$ | 37.35 ± 0.33 | Genotype | 16.15 | 0.0006* | a |
| LS (7) | *mdm* $Iso_{Short}$ | 39.01 ± 0.55 | Condition | 14.92 | <0.0001* | b |
| LS (17) | WT $Iso_{Long}$ | 35.88 ± 0.39 | Interaction | 3.80 | 0.032* | c |
| LS (7) | *mdm* $Iso_{Long}$ | 37.33 ± 0.58 | | | | a |
| LS (10) | WT $Iso_{RFE}$ | 34.12 ± 0.75 | | | | d |
| LS (7) | *mdm* $Iso_{RFE}$ | 37.77 ± 0.44 | | | | a, b |
| $\sigma_A$ (8) | WT $Iso_{Short}$ | 0.17 ± 0.01 | Genotype | 9.57 | 0.007* | a |
| $\sigma_A$ (5) | *mdm* $Iso_{Short}$ | 0.22 ± 0.02 | Condition | 37.79 | <0.0001* | b |
| $\sigma_A$ (12) | WT $Iso_{Long}$ | 0.12 ± 0.01 | Interaction | 0.77 | 0.47 | b |
| $\sigma_A$ (5) | *mdm* $Iso_{Long}$ | 0.15 ± 0.004 | | | | |
| $\sigma_A$ (7) | WT $Iso_{RFE}$ | 0.12 ± 0.01 | | | | |
| $\sigma_A$ (6) | *mdm* $Iso_{RFE}$ | 0.16 ± 0.02 | | | | |
| $I_{1,1}/I_{1,0}$ (10) | WT $Iso_{Short}$ | 1.50 ± 0.45 | Genotype | 39.80 | <0.0001* | a |
| $I_{1,1}/I_{1,0}$ (7) | *mdm* $Iso_{Short}$ | 9.12 ± 2.10 | Condition | 1.99 | 0.1535 | b |
| $I_{1,1}/I_{1,0}$ (16) | WT $Iso_{Long}$ | 1.43 ± 0.22 | Interaction | 2.27 | 0.1203 | a |
| $I_{1,1}/I_{1,0}$ (7) | *mdm* $Iso_{Long}$ | 5.96 ± 1.16 | | | | b |
| $I_{1,1}/I_{1,0}$ (8) | WT $Iso_{RFE}$ | 1.35 ± 0.45 | | | | a |
| $I_{1,1}/I_{1,0}$ (7) | *mdm* $Iso_{RFE}$ | 9.35 ± 1.53 | | | | b |
| $S_{M3}$ (12) | WT $Iso_{Short}$ | 14.29 ± 0.02 | Genotype | 2.35 | 0.1413 | a |
| $S_{M3}$ (7) | *mdm* $Iso_{Short}$ | 14.32 ± 0.01 | Condition | 12.45 | <.0001 | b |
| $S_{M3}$ (18) | WT $Iso_{Long}$ | 14.36 ± 0.02 | Interaction | 0.32 | 0.7297 | b |
| $S_{M3}$ (7) | *mdm* $Iso_{Long}$ | 14.38 ± 0.03 | | | | |
| $S_{M3}$ (12) | WT $Iso_{RFE}$ | 14.36 ± 0.02 | | | | |
| $S_{M3}$ (7) | *mdm* $Iso_{RFE}$ | 14.40 ± 0.01 | | | | |
| $S_{M6}$ (9) | WT $Iso_{Short}$ | 7.25 ± 0.006 | Genotype | 16.17 | 0.0007* | a, b |
| $S_{M6}$ (7) | *mdm* $Iso_{Short}$ | 7.24 ± 0.006 | Condition | 34.92 | <0.0001* | a |
| $S_{M6}$ (16) | WT $Iso_{Long}$ | 7.27 ± 0.004 | Interaction | 8.29 | 0.0015* | b |
| $S_{M6}$ (7) | *mdm* $Iso_{Long}$ | 7.25 ± 0.003 | | | | c |
| $S_{M6}$ (7) | WT $Iso_{RFE}$ | 7.30 ± 0.004 | | | | d |
| $S_{M6}$ (6) | *mdm* $Iso_{RFE}$ | 7.26 ± 0.003 | | | | b |
| $S_{A6}$ (9) | WT $Iso_{Short}$ | 5.84 ± 0.003 | Genotype | 10.27 | 0.007* | a |
| $S_{A6}$ (4) | *mdm* $Iso_{Short}$ | 5.84 ± 0.009 | Condition | 36.09 | <0.0001* | a |
| $S_{A6}$ (9) | WT $Iso_{Long}$ | 5.90 ± 0.005 | Interaction | 3.97 | 0.03* | b |
| $S_{A6}$ (5) | *mdm* $Iso_{Long}$ | 5.88 0.002 | | | | c, d |
| $S_{A6}$ (7) | WT $Iso_{RFE}$ | 5.89 ± 0.007 | | | | b, c |
| $S_{A6}$ (5) | *mdm* $Iso_{RFE}$ | 5.85 ± 0.002 | | | | a, d |

**Data S1. Source_Data (Separate Excel file)**
